# Structure and enzymology of glutaminase mutants that disrupt glutamine-glutamate homeostasis and cause neurological disease

**DOI:** 10.1101/2025.09.14.674662

**Authors:** Cléa S. Crane, Thora K. McIssac, Shawn K. Milano, Richard A. Cerione, Scott M. Ulrich

## Abstract

The glutaminase (GLS) isoforms KGA and GAC are expressed in neurons where they hydrolyze glutamine to produce the excitatory neurotransmitter glutamate. Two *de novo* gain-of-function mutants of GLS, S482C and H461L, were recently identified in patients with developmental delay, epilepsy, and infantile cataract. These patients exhibited high glutamate and low glutamine concentrations in the brain, suggesting that the GLS mutants have abnormal enzymology. Here, we examined the enzymatic properties of these GLS mutants and found that they exhibit a total (S482C) or partial (H461L) loss of glutamate product inhibition, lifting this restriction on glutamate accumulation. The mutant enzymes also no longer require the anionic activator phosphate to stimulate enzymatic activity or induce filament formation. Structural analysis of the S482C GAC mutant shows the mutation shifts the key catalytic residue Y466 into the catalytically competent position and disrupts a key hydrogen bond between it and the glutamate product, explaining how the S482C mutant has enzymatic activity in the absence of phosphate and is insensitive to glutamate product inhibition. These results shed new light on the mechanism of phosphate activation and glutamate product inhibition of GLS and show that loss of these enzymatic properties disrupts glutamate homeostasis in the brain and causes neurological disease.

## Introduction

Glutaminases are tetrameric mitochondrial enzymes that catalyze the hydrolysis of glutamine to glutamate. There are two glutaminase genes in humans; *GLS* encodes kidney-type glutaminase (KGA) and a C-terminal splice variant glutaminase C (GAC), and *GLS2* encodes liver-type glutaminase (LGA).^1,2^ *GLS* isoforms KGA and GAC (collectively referred to here as GLS) are highly expressed in the brain and are the main producers of the excitatory neurotransmitter glutamate. Glutamate homeostasis in the brain is mediated by GLS and the opposing enzyme glutamine synthetase (GS) in the glutamine-glutamate cycle.^3^ Presynaptic neurons release glutamate to mediate excitatory signals in postsynaptic neurons. Glutamate is cleared from the synaptic cleft into astrocytes by high-affinity transporters, where it is converted to glutamine by GS. Glutamine is then released from astrocytes and taken into neurons where it is hydrolyzed to glutamate by GLS to complete the cycle.^4,5^

Aberrant GLS activity can disrupt glutamine/glutamate homeostasis in the brain and cause neurological disease. Loss-of-function GLS mutations cause ataxia, neonatal seizures, cerebral edema and structural brain abnormalities.^6,7^ Elevated GLS activity in the brain causes excess glutamate and excitotoxicity, a pathology of glutamatergic neurons that is a component of many neurological diseases including Alzheimer’s disease, amyotrophic lateral sclerosis (ALS), and epilepsy.^8–13^ HIV-infected microglia upregulate GLS resulting in excess brain glutamate, which contributes to HIV-associated neurocognitive disorder (HAND).^14^ Similarly, transgenic mice engineered to overexpress the GAC isoform in the brain have excess glutamate and display learning defects and synaptic dysfunction.^15^

Recent reports identified *de novo* gain-of-function GLS mutations in two patients with neurological disease. One patient carried a S482C mutation and suffered from infantile cataract, skin lesions, and profound developmental delay.^16^ The second patient carried a H461L mutation and suffered from epilepsy and moderate developmental delay.^17^ Both patients exhibit low glutamine and high glutamate levels in the brain, which is the probable cause of their disease.^18^ The S482C and H461L mutations reside in the catalytic domain of GLS, which is identical in the KGA and GAC isoforms. S482 is adjacent to the active site and contacts Y466, a key catalytic residue. H461 is a surface residue in the monomer-monomer interface where it contacts its twin on the adjoining subunit (Figure 1). Sequence alignment shows that S482 is strictly conserved across the glutaminase superfamily while H461 shows some variation (Figure S1).

**Figure 1.**
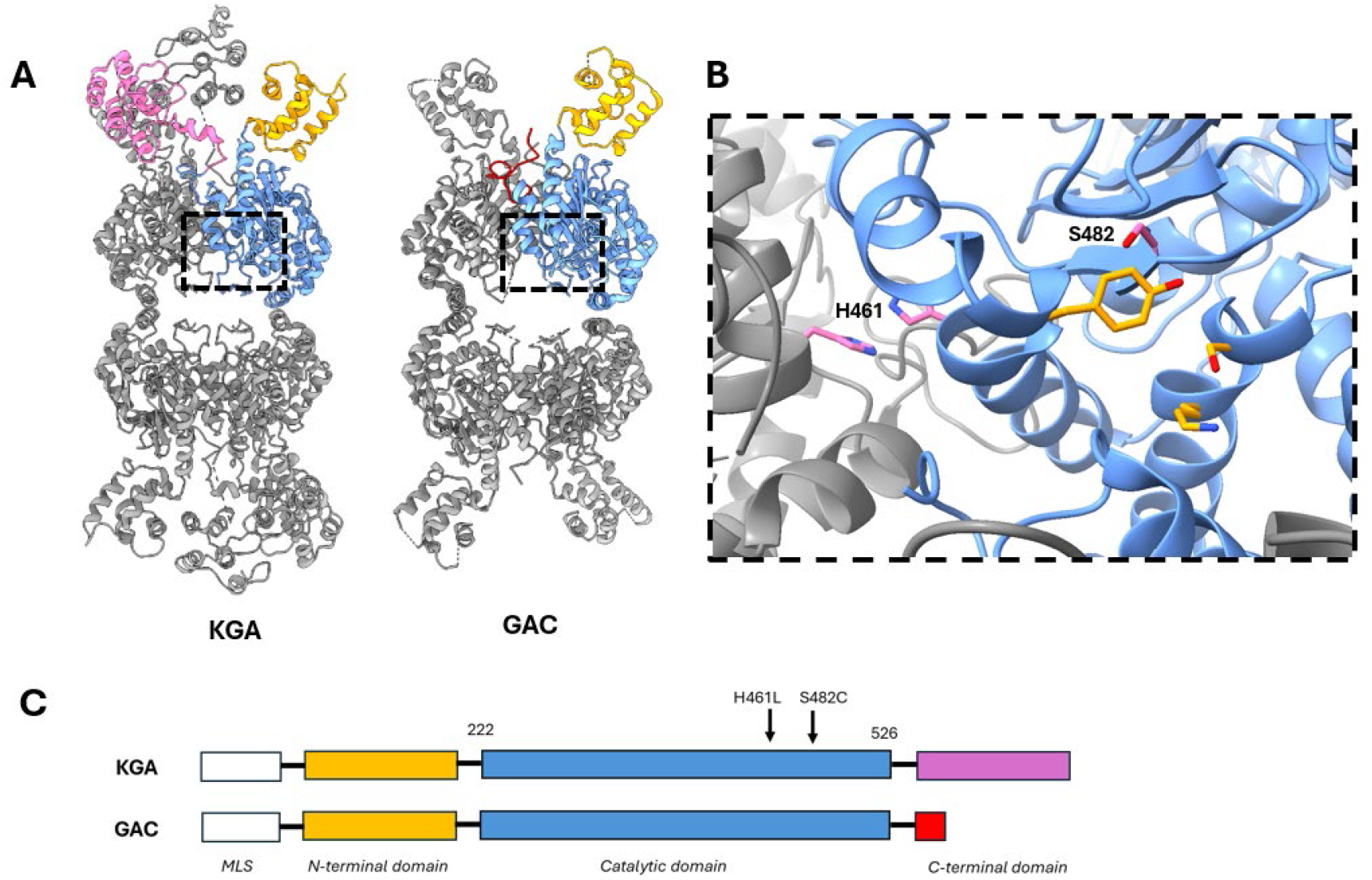
Structure of the GLS isoforms and the disease-associated mutations. **A)** Structure of GLS isoforms KGA and GAC. The isoforms have identical N-terminal (orange) and catalytic (blue) domains and different C-terminal domains. **B)** Detailed view of the GLS active site showing key amino acids involved in the catalytic mechanism: S286, K289, and Y466 (orange). The residues mutated in the patients are S482 and H461 (pink). S482 contacts catalytic residue Y466 and H461 is at the monomer-monomer interface where it contacts its twin across the interface.C)Domain structure of KGA and GAC showing the positions of the disease-associated mutations in the shared catalytic domain. Images generated from PDB IDs 5UQE (KGA) and 3UNW (GAC).

The reports identifying the S482C and H461L GLS mutations proposed that the glutamate excess in the patients was due to hyperactivity of the mutant GLS enzymes, but their enzymology was not studied in detail. Here, we sought to characterize the enzymatic properties of the S482C and H461L GLS mutants. KGA and GAC are complex enzymes; their catalytic activity is regulated by multiple ligands that also affect their ability to reversibly assemble into filament-like structures. Understanding how the S482C and H461L mutations alter the enzymatic properties of GLS could shed new light on GLS enzymology, explain the excess glutamate observed in the patients, and identify the biochemical properties of GLS that are important to maintain glutamate homeostasis in the brain.

## Results and Discussion

### S482C and H461L GLS mutants do not require phosphate for enzymatic activity

Inorganic phosphate is an essential activator of GLS enzymes.^1,19^ We measured the effect of phosphate on the activity of wild-type (WT) KGA and GAC as well as the S482C and H461L mutants. We found that WT KGA and GAC exhibited no enzymatic activity in the absence of phosphate and are strongly activated by it, with EC_50_ values of 30 ± 4 mM (KGA) and 16 ± 2 mM (GAC), consistent with previous reports.^19,20^ Conversely, the S482C and H461L mutants of both GLS isoforms are active enzymes in the absence of phosphate and show little or no stimulation upon phosphate addition (Figure 2A). The GLS isoforms localize to mitochondria where the phosphate concentration is 15 mM,^21,22^ sufficient to activate the enzymes. In neurons, BCH domain family members BNIP-H/Caytaxin^23^ and BMCC1S^24^ have been reported to transport KGA to the cytoplasm, where the phosphate concentration is significantly lower.^21^ The report showed that overexpression of BNIP-H/Caytaxin in 293T cells lowered glutamate levels, consistent with deactivation of KGA by movement to a lower phosphate environment.^23^ The loss of phosphate activation by the S482C and H461L GLS mutants would result in elevated enzymatic activity under the low phosphate conditions in the cytoplasm and thus could contribute to the excess glutamate observed in the patients.

**Figure 2:**
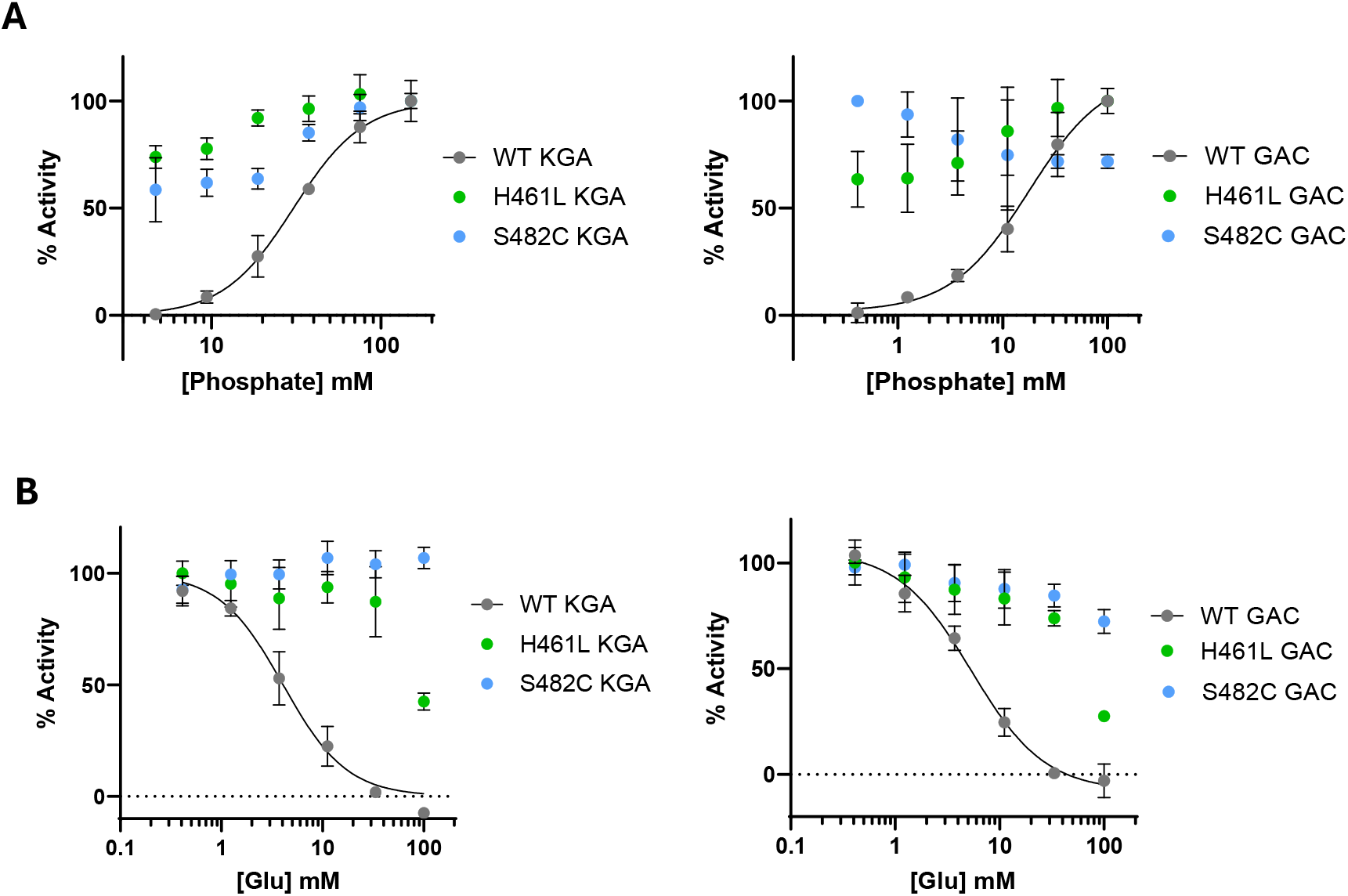
Phosphate activation and glutamate product inhibition of WT KGA and GAC and the S482C and H461L mutants. **A)** Plots of the enzymatic activity of WT KGA and GAC and the S482C and H461L mutants with increasing phosphate concentration, normalized to the maximal signal. **B)** Plots of the enzymatic activity of WT KGA and GAC and the S482C and H461L mutants with increasing glutamate concentration. The phosphate and glutamine concentrations were fixed at 15 mM each. *Data are mean ± S*.*D*., *n = 3*.

### S482C and H461L GLS mutants are resistant to glutamate product inhibition

GLS enzymes exhibit product inhibition by glutamate, which binds the active site and is competitive with phosphate.^1,25,26^ We measured glutamate product inhibition of WT KGA and GAC as well as the S482C and H461L mutants at 15 mM glutamine, about 5-fold above its K_0.5_ value for WT enzymes, and at 15 mM phosphate, equivalent to its concentration in mitochondria. Under these conditions, we found that WT KGA and GAC both exhibit product inhibition by glutamate with IC_50_ values of 4.1 ± 0.5 mM (KGA) and 5.3 ± 0.7 mM (GAC), whereas the mutants showed a complete (S482C) and partial (H461L) loss of inhibition by glutamate (Figure 2B). The glutamate concentration in neuronal mitochondria (as well as whole brain tissue) is reported to be 10 mM, ^27–29^ which is sufficient to inhibit WT KGA and GAC and thereby suppress the accumulation of excess glutamate. The S482C and H461L mutants are active enzymes at 10 mM glutamate and can produce glutamate above that level. It is likely that glutamate product inhibition is an important property of GLS that imposes an upper limit on glutamate concentration in the brain and the loss of product inhibition by the S482C and H461L mutants contributes to the excess glutamate observed in the patients.

### The S482C and H461L mutations have modest effects on enzyme kinetics

We measured the k_cat_ and K_0.5_ values of WT KGA and GAC as well as the S482C and H461L mutants to determine if the mutants have enhanced enzymatic activity relative to the wild-type enzymes. We chose to conduct these measurements at 50 mM phosphate for all enzymes, which is twice the EC_50_ for KGA activation and three times the EC_50_ for GAC activation. Under these conditions, we found that the S482C and H461L mutants of KGA and GAC have comparable kinetic parameters to the corresponding phosphate-activated WT enzymes (Figure 3 and Table 1). Thus, the excess glutamate observed in the patients is not due to increased catalytic capacity of the mutant enzymes but is instead due to the loss of phosphate activation and/or glutamate product inhibition.

**Table 1.** Kinetic parameters of WT KGA and GAC and the S482C and H461L mutants. *Data are best-fit values ± SEM*

| Enzyme | $k_{\text{cat}}$ ($\text{s}^{-1}$) | $K_{0.5}$ glutamine<br>(mM) | Hill slope | $\text{EC}_{50}$ phosphate<br>(mM) | $\text{IC}_{50}$ glutamate<br>(mM) |
| --- | --- | --- | --- | --- | --- |
| WT KGA | $7.0 \pm 0.5$ | $3.8 \pm 0.4$ | $1.2 \pm 0.1$ | $30 \pm 4$ | $4.1 \pm 0.5$ |
| S482C KGA | $6.6 \pm 0.5$ | $10.1 \pm 0.7$ | $1.9 \pm 0.1$ | - | - |
| H461L KGA | $3.9 \pm 0.8$ | $1.7 \pm 0.3$ | $1.1 \pm 0.2$ | - | - |
| WT GAC | $16 \pm 1$ | $2.9 \pm 0.3$ | $1.1 \pm 0.1$ | $16 \pm 2$ | $5.3 \pm 0.7$ |
| S482C GAC | $13.1 \pm 0.9$ | $5.1 \pm 0.3$ | $1.9 \pm 0.1$ | - | - |
| H461L GAC | $13.4 \pm 0.8$ | $2.0 \pm 0.2$ | $1.1 \pm 0.1$ | - | - |

**Figure 3.**
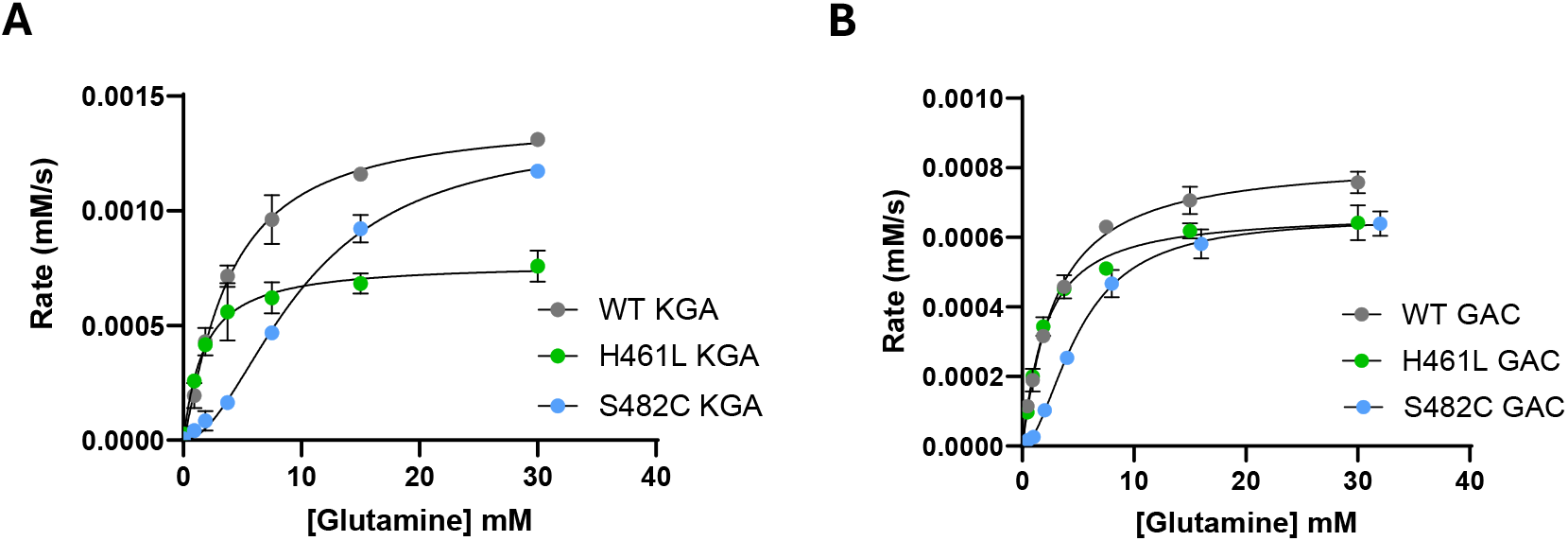
Glutamine substrate concentration vs reaction rate plots. **A)** WT KGA and the S482C and H461L mutants. **B)** WT GAC and the S482C and H461L mutants. All reactions were carried out in the presence of 50 mM phosphate. *Data are mean ± S*.*D*., *n = 3*.

Surprisingly, while the substrate-velocity curves for WT KGA, WT GAC, and the H461L mutants are hyperbolic, consistent with previous reports,^20,30,31^ the S482C mutants of both KGA and GAC exhibit sigmoidal kinetics, indicating a shift to positive substrate cooperativity (Figure 3). The LGA enzyme encoded by the *GLS2* gene also shows substrate cooperativity, no glutamate product inhibition, and weak activation by phosphate.^1,20,31,32^ The catalytic domains of LGA and the GLS isoforms are 80% identical, and the cause of their different enzymatic properties remains unclear.^2,33^ The observation that the S482C mutation in GLS results in LGA-like enzymatic properties could help address this question.

### The S482C and H461L mutants constitutively assemble into filaments

Glutaminase enzymes are among a growing number of metabolic enzymes that reversibly assemble into oligomeric filament-like structures.^34,35^ GLS filaments are the active form of the enzyme; phosphate induces filament formation while glutamate causes the filaments to disassemble into tetramers.^36–39^ Since we observed that the S482C and H461L mutants do not require phosphate for enzymatic activity and are resistant to glutamate product inhibition, we suspected that the regulation of filament formation by the mutants would also be altered. A recent report on GLS filamentation included an analysis of S482C GAC, which was shown by electron microscopy to form filaments in the absence of phosphate.^40^ This report also showed that dimethyl glutamate (a cell-permeable glutamate precursor) caused dissociation of WT GAC filaments but not S482C GAC filaments in cells. We evaluated filament formation by S482C and H461L GAC in the absence of phosphate using negative-stain electron microscopy. Our data show that the S482C mutant forms filaments in the absence of phosphate, in agreement with this previous report. We found that H461L GAC also forms filaments in the absence of phosphate, although the filaments appear shorter than S482C GAC (Figure 4A). These results align with our observation that the S482C and H461L GLS mutants are active enzymes in the absence of phosphate.

**Figure 4.**
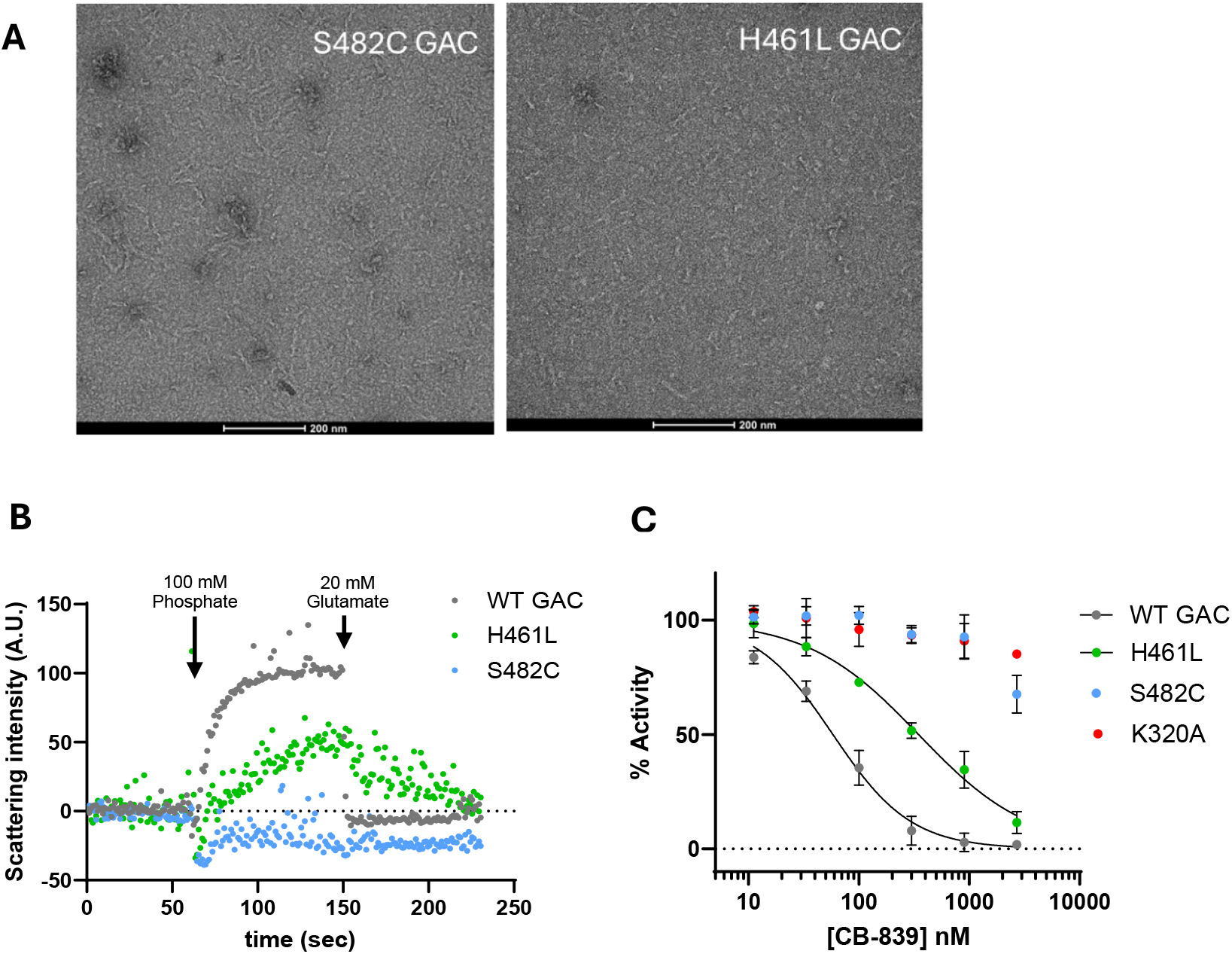
Filament formation by WT GAC and the H461L and S482C mutants. **A)** Negative-stain electron microscopy images of S482C and H461L GAC show that both mutants form filaments in the absence of phosphate. **B)** Right-angle light scattering (RALS) assay showing real-time filamentation of WT GAC and the S482C and H461L mutants in response to phosphate and glutamate. **C)** Dose-response curves of the GLS inhibitor CB839 on the enzymatic activity of WT GAC and the H461L, S482C and K320A GAC mutants. *Data are mean ± S*.*D*., *n = 3*.

We measured filament formation of WT GAC and the S482C and H461L mutants using right-angle light scattering (RALS) as a real-time assay.^38,41^ We confirmed that phosphate promotes filament formation of WT GAC, indicated by an increase in light scattering upon phosphate addition. Subsequent addition of glutamate dissociates WT GAC filaments, shown by a return to the basal level of light scattering (Figure 4B). Conversely, the S482C mutant showed no change in light scattering upon the addition of phosphate or glutamate, consistent with constitutive filament formation. The H461L mutant showed a slight increase in light scattering upon phosphate addition which was reversed by subsequent addition of glutamate, suggesting that phosphate can lengthen or enhance H461L GAC filaments and glutamate counters this effect. These results align with our observation that the S482C and H461L GLS mutants are insensitive (S482C) or resistant (H461L) to glutamate product inhibition.

We also tested whether the S482C and H461L GAC mutants are sensitive to the GLS inhibitor CB839. CB839 binds at the dimer-dimer interface of GLS, which is remote from the S482C and H461L mutations (Figure S2).^42^ CB839 binding locks GLS into inactive tetramers, and conditions that promote GLS filament formation cause resistance to CB839. For example, the K320A GLS mutant is a constitutive filament and is resistant to CB839, as is WT GLS under high phosphate concentrations.^43–45^ As such, CB839 sensitivity can be used as a proxy measurement for GLS filamentation. We found that CB839 potently inhibited WT GAC (IC_50_ = 58 *±* 4 nM) while the K320A mutant was insensitive to it, as expected (Figure 4C). The S482C GAC mutant was also insensitive to CB839, consistent with constitutive filament formation. CB839 inhibited the H461L mutant (IC_50_ = 350 *±* 30 nM) but to a lesser degree than WT GAC. Together, these results show that both the S482C and H461L mutants are constitutive filaments and suggest that H461L GAC forms short and/or weak filaments compared to S482C GAC.

### X-ray crystal structure of S482C GAC

We solved the X-ray crystal structure of the S482C GAC mutant (PDB ID: 9PIA) to understand how the mutation causes the loss of phosphate activation and glutamate product inhibition. We found that the only significant differences between the structure of the S482C mutant and apo WT GAC are the positions of the mutant S482C residue and Y466, a key catalytic residue. This was not unexpected since the S482 hydroxyl group contacts the edge of the Y466 ring in the apo WT GAC structure.^46^ In the S482C GAC structure, the mutant cysteine residue rotates away from Y466 to pack against the neighboring hydrophobic residue M465, both of which reside in the turn of a short helix-turn-helix motif. The packing of the mutant cysteine against M465 induces a small dip in this elbow-like motif which repositions Y466 (Figure 5A).

**Figure 5.**
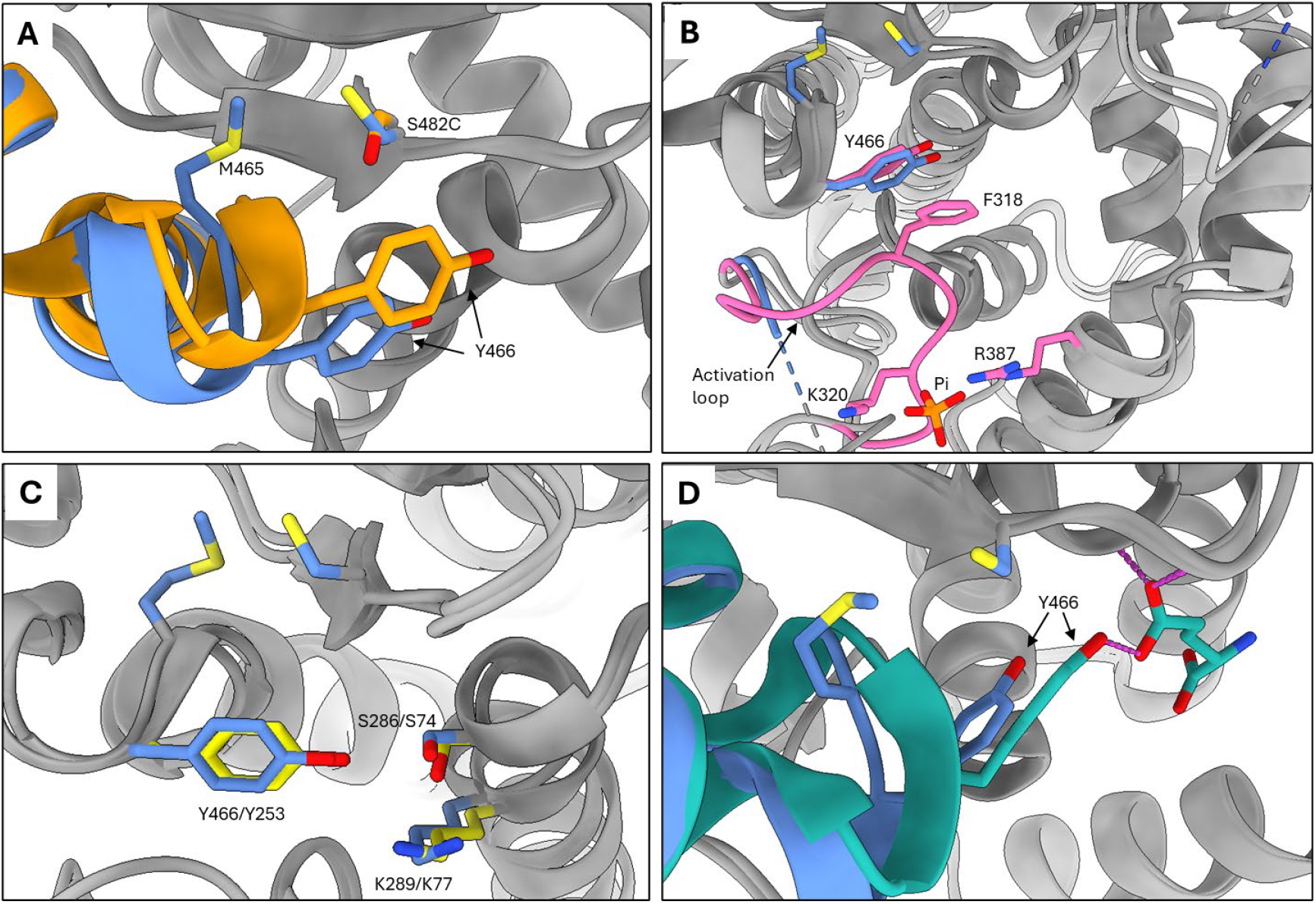
X-ray crystal structure of the S482C GAC mutant. **A)** Superimposed structures of inactive apo WT GAC (orange) and active S482C GAC (blue). S482 of WT GAC contacts the catalytic residue Y466. Cysteine 482 of GAC S482C is rotated away from Y466 to pack against M465, causing a small displacement of the helix-turn-helix motif and shifting Y466 to a “down and in” configuration. **B)** Superimposed structures of active S482C GAC (blue) and active phosphate-bound WT GAC (pink) show the same Y466 position. Phosphate promotes this configuration by ordering the activation loop which promotes a Π-stacking interaction between F318 and Y466. **C)** Superimposed structures of the S482C GAC (blue) and *B. subtilis* glutaminase YbgJ (yellow). Shown are the Ser-Lys-Tyr catalytic triad of each enzyme, indicating that the position of Y466 in S482C GAC is the catalytically competent configuration. **D)** Superimposed structures of active S482C GAC (blue) and inhibited glutamate-bound WT GAC (teal). The Y466 hydroxyl group makes a weak hydrogen bond to the glutamate side chain. The configuration of Y466 induced by the S482C mutation is out of hydrogen bond range. Images generated from PDB IDs 9PIA (this study, S482C GAC); 3VOY (apo WT GAC); 8IMA (phosphate-bound WT GAC); 3UNW (glutamate-bound WT GAC); 1MKI (*B. subtilis* YbgJ) superimposed using the USCF ChimeraX Needleman–Wunsch tool for best aligning chains.^51^

The repositioning of Y466 by the S482C mutation explains how the S482C GAC mutant is active in the absence of phosphate. Phosphate binds GLS near the dimer-dimer interface using K320 and R387 as well as Y394’ and K398’ from the subunit across the interface. K320 is part of the GLS activation loop (L^316^RFNKLF^322^), which forms the rim of the active site and is disordered in the inactive apo WT enzyme. Phosphate binding causes the activation loop to become ordered, which initiates a Π-stacking interaction between activation loop residue F318 and Y466. This interaction repositions Y466, allowing it to participate in a proton relay system with two other catalytic residues S286 and K289, accounting for the observed phosphate stimulation of enzymatic activity.^37–39^ This mechanism is supported by the observation that the F318S GAC mutant, which cannot engage in the Π-stacking interaction with Y466, is inactive and does not respond to phosphate.^37^ Superimposing the structures of active phosphate-bound WT GAC and the S482C mutant shows that the position of Y466 in the S482C structure is identical to its position in the phosphate-bound WT structure. The activation loop in the S482C mutant remains disordered, indicating that phosphate binding to the WT enzyme and the S482C mutation promote the same catalytically active configuration of Y466 but in mechanistically distinct ways (Figure 5B).

Comparing the structures of the S482C GAC mutant and *B. subtilis* glutaminase YbgJ supports the idea that the position of Y466 in the S482C GAC structure is the active configuration. The catalytic domain of bacterial and mammalian glutaminases share a β-lactamase fold and use a common catalytic triad consisting of S286, K289, and Y466 in GAC and S74, K77, and Y253 in YbgJ.^47^ YbgJ is not stimulated by phosphate nor inhibited by the glutamate product, so the configuration of the catalytic triad in the YbgJ structure very likely represents the active form of the enzyme. The configuration of the catalytic triad of the S482C GAC mutant is nearly identical to that of YbgJ, suggesting that the catalytic triad of the S482C mutant is in its catalytically active configuration (Figure 5C).

The repositioning of Y466 by the S482C mutation can also account for the loss of glutamate product inhibition by the mutant enzyme. The structure of WT GAC with glutamate bound to the active site shows the hydroxyl group of Y466 makes a weak hydrogen bond to the glutamate side-chain carboxylate, with a donor-acceptor distance of 3.3 Å.^42^ Superimposing the structures of S482C GAC and glutamate-bound WT GAC shows that the position of Y466 imposed by the S482C mutation extends the donor-acceptor distance to 3.9 Å. This is out of hydrogen-bond range and can explain the S482C GAC mutant enzyme’s lower affinity for glutamate and the loss of product inhibition (Figure 5D).

Together, these results show that there are two key positions of Y466. One position is catalytically active and has a low affinity for glutamate, as seen in the structures of phosphate-bound WT GAC, S482C GAC, and bacterial glutaminases. The second position is catalytically inactive and has a higher affinity for glutamate, as seen in the structures of apo and glutamate-bound WT GAC.

## Conclusion

In this study, we examined the enzymatic properties of the S482C and H461L GLS mutants discovered in two patients that exhibit high concentrations of glutamate in the brain, neurological disease, and developmental delay. We found that both mutant enzymes are active in the absence of phosphate, a required activator of WT GLS. Similarly, neither mutant enzyme requires phosphate to assemble into filaments, which is the active form of the enzyme. The mutants also show a complete (S482C) or partial (H461L) loss of product inhibition by glutamate. Both mutants exhibit k_cat_ and K_0.5_ values comparable to those of the phosphate-activated wild-type enzymes. Therefore, it is the loss of phosphate activation and/or glutamate product inhibition that accounts for the excess glutamate and neurological disease observed in the patients. The structure of the S482C GAC mutant shows that the catalytic residue Y466 shifts to its catalytically competent configuration and disrupts a hydrogen bond between the Y466 hydroxyl group and the glutamate product, explaining the activity of the mutant enzyme in the absence of phosphate and the loss of glutamate product inhibition. These results support a Y466-centric mechanism of phosphate activation and glutamate product inhibition of GLS enzymatic activity.

## Data availability

The model and electron density map of GAC (S482C) are available from the Protein Data Bank under PDB ID 9PIA. All other data are contained within the manuscript and the supporting information section.

## Acknowledgements

The authors would like to thank Dr. Shi Feng for helpful discussions and help with the RALS assay, Biswanath Shaw and Saul De La Pena for help with negative stain electron microscopy, and the Crane lab for the generous use of instrumentation. This study was supported by a grant from the NIH (R35 GM152206 to R.A.C.). The collection of negative stain images relied on using an instrument supported by NIH award S10OD030470-01. Collection of X-ray crystallography data was done at the Macromolecular Diffraction at CHESS (MacCHESS) facility, which is supported by NIH NIGMS award 1-P30-GM124166.

## Author contributions

SMU, SKM, RAC: Design and supervision of the study; CSC, TKM, SKM, SMU: Data collection and analysis; All authors: Drafting and revising the manuscript; RAC: Funding acquisition.

**Correspondence** and requests for materials should be addressed to Richard A. Cerione.

## Experimental methods

### Recombinant glutaminase expression and purification

An N-terminal 6His-tagged form of human KGA without the mitochondrial localization sequence (residues 1-71) was cloned into the pET28a plasmid. An N-terminal 6His-tagged form of human GAC without the mitochondrial localization sequence (residues 1-71) was cloned into the pQE80L plasmid.^20^ Site-directed mutagenesis was performed using Phusion DNA polymerase (New England Biolabs). The primers (5′–3′) used to introduce the mutations in GAC and KGA were as follows:

S482C: CTT CCT GCA AAA TGT GGA GTT GCT GGG (forward); CCC AGC AAC TCC ACA TTT TGC AGG AAG (reverse)

H461L: TTG AGT TTG ATG CTT TCC TGT GGC ATG (forward); CAT GCC ACA GGA AAG CAT CAA ACT CAA (reverse)

K320A: CTA AGA TTC AAC GCA CTA TTT TTG AAT (forward); ATT CAA AAA TAG TGC GTT GAA TCT TAG (reverse)

Expression and purification of glutaminase enzymes was carried out by transforming the constructs described above into *E. coli* BL21(DE3) competent cells (New England Biolabs), which were then grown in LB media overnight with 50 μg/mL kanamycin (KGA) or 50 μg/mL ampicillin (GAC). The starter cultures were used to inoculate 6 L of terrific broth (1:100 dilution) with the same antibiotic concentrations and shaken at 37 °C, 180 rpm for 3–4 h until the OD_600_ reached between 0.6 and 0.8. The flasks were chilled at 4 °C for 1–2 h before induction with 100 μM IPTG and shaken at room temperature (RT) at 180 rpm for 16 h. Cells were collected by centrifugation at 5000 × g for 10 minutes and frozen. Frozen cell pellets were resuspended in 150 mL lysis buffer (50 mM Tris–HCl pH 8.5, 500 mM NaCl, 10% glycerol) supplemented with protease inhibitor cocktail (Roche) then lysed by sonication. The mixture was clarified by ultra-centrifugation (40,000 × g) for 45 min. The supernatant was then loaded onto Co^2+^ charged TALON resin (GoldBio), previously equilibrated with wash buffer (50 mM Tris–HCl pH 8.5, 10 mM NaCl, 10 mM imidazole). The protein that bound to the column was washed with wash buffer (150 mL) and eluted with wash buffer supplemented with 320 mM imidazole. Further purification was performed by anion exchange chromatography using HiTrap Q HP column (Cytiva) and size-exclusion chromatography using Superdex 200 pg 16/600 column (GE Healthcare). Proteins were kept in 20 mM Tris–HCl pH 8.5, 150 mM NaCl, snap-frozen in liquid nitrogen, and stored at −80 °C. Protein concentrations were determined by absorbance at 280 nm using extinction coefficients calculated using the Expasy ProtParam tool.

### Glutaminase enzyme assays

Glutaminase was diluted in glutaminase buffer (65 mM Tris acetate pH 8.6, 0.2 mM EDTA) to a final concentration of 50 nM (GAC) or 200 nM (KGA) for each type of assay described below.

#### Michaelis-Menten kinetics

For the measurement of Michelis-Menten kinetics, 80 µL of the enzyme mixture was added to 96 well plates. The glutaminase reaction was initiated by the addition of 20 µL of a solution of glutamine (150, 75, 37.5, 18.75, 9.37, 4.69 mM) and K_2_HPO_4_ (250 mM) in glutaminase buffer, mixed by gently pipetting up and down and incubated at room temperature for 600 seconds.

#### Phosphate stimulation

The assay for phosphate stimulation was similar except the reaction was initiated by the addition of 20 µL of a solution of glutamine (100 mM) and K_2_HPO_4_ (500, 250, 125, 62.5, 31.2, 15.6, 7.8, 0 mM) in glutaminase buffer, mixed by gently pipetting up and down and incubated at room temperature for 600 seconds.

#### CB839 inhibition

The assay for CB-839 inhibition was similar except the enzyme mixture was incubated with 1.0 µL of a DMSO solution of CB839 and mixed by gently pipetting up and down. The reaction was initiated by the addition of 20 µL of a solution of glutamine (100 mM) and K_2_HPO_4_ (500 mM) in glutaminase buffer and incubated at room temperature for 600 seconds.

In the assays described above, the glutaminase reactions were quenched by the addition of 10 µL of cold HCl (3 M). An aliquot (10 µL) of each quenched glutaminase reaction was added to 190 µL of a glutamate dehydrogenase reaction, which consisted of Tris-HCl (100 mM, pH 9.4), NAD+ (2 mM), glutamate dehydrogenase (2 µL of a 50% glycerol solution, ≥35 units/mg protein), and hydrazine (1 µL) then incubated at room temperature for 40 min. The absorbance increase at 340 nm was measured and converted to glutamate concentrations using the extinction coefficient for NADH (6220 M^-1^ cm^-1^).

#### Glutamate inhibition

The assay to measure glutamate product inhibition used a smaller initial enzyme volume (70 µL) to which was added a solution of glutamate (10 µL) in glutaminase assay buffer (1000 mM, 333 mM, 111 mM, 37.0 mM, 12.3 mM, 4.1 mM, 0 mM). The reaction was initiated by the addition of 20 µL of a solution of glutamine (75 mM) and K_2_HPO_4_ (75 mM) in glutaminase buffer such that the final concentration of each was 15 mM, mixed by gently pipetting up and down and incubated at room temperature for 600 seconds. The reactions were quenched by addition of 10 µL of HCl (3 M). An aliquot (10 µL) of each quenched glutaminase reaction was added to 190 µL of a glutamate dehydrogenase reaction configured to measure the ammonia product which consisted of Tris-HCl (100 mM, pH 9.4), NADH (0.23 mM), α-ketoglutarate (3.4 mM), and glutamate dehydrogenase (2 µL of a 50% glycerol solution, ≥35 units/mg protein) then incubated at room temperature for 40 min. The decrease in absorbance at 340 nm was measured and converted to ammonia concentrations using the extinction coefficient for NADH (6220 M^-1^ cm^-1^). Kinetic parameters, EC_50_ and IC_50_ values were determined by using the appropriate curve fitting equation in the GraphPad Prism software (GraphPad Software, San Diego, CA).

### Negative stain electron microscopy

Formvar/carbon film 200 mesh copper grids (Electron Microscopy Sciences, EMS) were plasma cleaned by using PELCO easiGlow system (TED PELLA). Glutaminase was dissolved in 20 mM Tris pH 8.5, 120 mM NaCl to a final concentration of 1 μM and inorganic phosphate K_2_HPO_4_ was added to a final concentration of 50 mM when needed. Ten microliters of the mixture were applied onto the grids for a 60 s incubation and the excess protein solution was blotted with filter paper. Ten microliters of 2% uranyl acetate was applied to the grid 30 sec staining followed by blotting two consecutive times. The grids were air dried for 5 min and visualized by Thermo Fisher F200Ci electron microscope at 120 keV.

### X-ray crystallography

Human GAC (S482C) was concentrated to 5 mg/mL using an Amicon ultrafiltration device (10 KDa cutoff; Millipore). Crystals were grown at 20 °C using the hanging drop vapor diffusion technique (2 μL of protein solution and 2 μL of reservoir solution), with the reservoir containing 10% PEG 6000 (w/v), 1 M LiCl, and 0.1 M Tris, pH 8.5. Crystals formed after 5 days. Glycerol was used as the cryoprotectant prior to plunge freezing. The diffraction data were collected at cryogenic temperature (100 K) at the Cornell High Energy Synchrotron Source (MacCHESS). The monomer extracted from the apo human WT GAC structure (Protein Data Bank ID: 5D3O) was used as the search model for molecular replacement. The data reduction was performed with HKL2000 prior to phasing and refinement using Phenix and Coot.^48–50^ The statistics of data collection and structure refinement are summarized in Table S1. The structure is deposited in the Protein data bank under PDB ID 9PIA.

### Right-angle light scattering (RALS)

Frozen glutaminase aliquots were thawed on ice and diluted to a final concentration of 2 μM in 1 mL buffer (20 mM Tris pH 8.5, 150 mM NaCl) and added in a 1.2 mL cuvette and inserted into a Varian Cary Eclipse fluorimeter. The cuvette temperature was set to 25 °C and stirred with a magnetic stir bar. The signal was recorded using excitation and emission wavelengths of 340 nm (5 nm bandpass). The samples were incubated until the signal was stable (~3-5 mins) and scattering intensity was recorded for 300 seconds during which time 20 μL of K_2_HPO_4_ (2.5 M) or glutamate (1 M) dissolved in the same buffer were added as indicated.

