## Supplemental information for "Structure and enzymology of glutaminase mutants that disrupt glutamine-glutamate homeostasis and cause neurological disease"

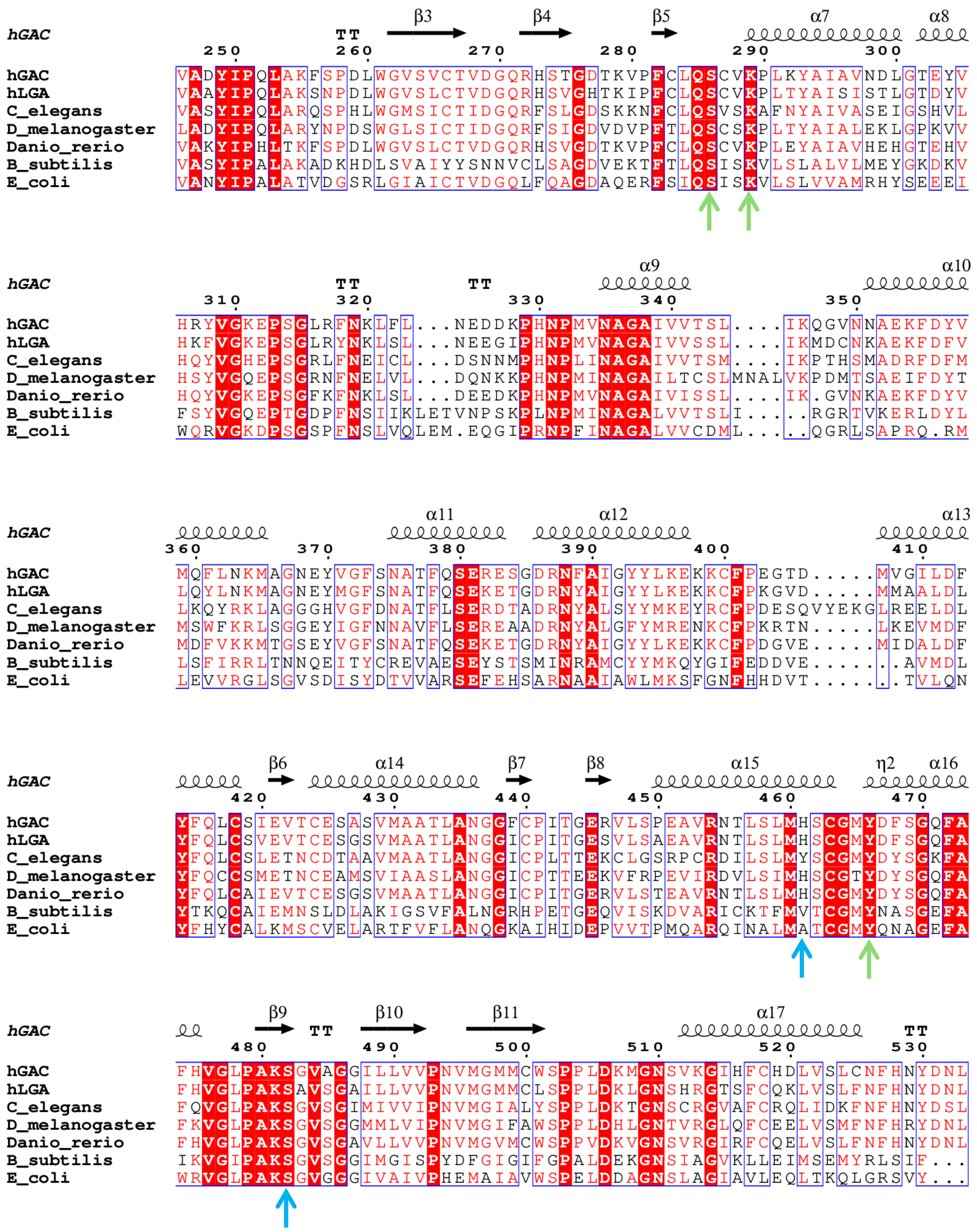


**Figure S1.** Sequence alignment of the catalytic domain of glutaminases including human GAC (the catalytic domain of KGA is identical to GAC), human LGA, and glutaminases from several model organisms. Amino acid numbering is for human GAC. The conserved catalytic residues S286, K289, and Y466 are marked with green arrows; the sites of the disease-associated mutations S482C and H461L are marked with light blue arrows.

**Table S1.** Crystallographic Data Collection and Model Refinement Statistics

| | |
|---|---|
| **Data Collection** | |
| Space group | P21 |
| Cell dimensions | |
| a (Å) | 50.85 |
| b (Å) | 138.21 |
| c (Å) | 177.12 |
| β (°) | 93.58 |
| Resolution (Å) | 50.0 - 3.0 |
| Unique reflections | 49294 (2404) |
| Redundancy | 6.9 (7.1) |
| Completeness (%) | 100.0 (100.0) |
| CC1/2 | 0.99 (0.52) |
| **Refinement** | |
| Resolution (Å) | 50 - 3.0 |
| $R_{work}/R_{free}$ | 0.19/0.21 |
| RMSD | |
| Bond length (Å) | 0.01 |
| Angle (°) | 1.08 |
| Ramachandran statistics | |
| Favored regions (%) | 94.96 |
| Allowed regions (%) | 4.46 |
| Outliers (%) | 0.57 |
| Avg B-factors (Å$^2$) | 71.93 |
| PDB ID | 9PIA |

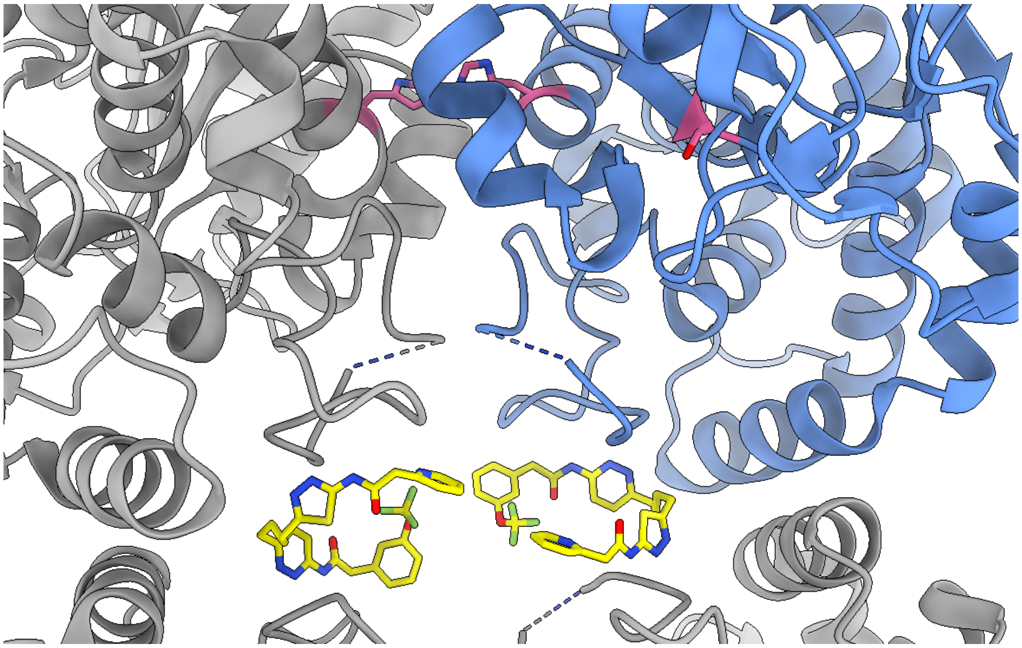

**Figure S2. Structure of CB839 bound to KGA.** CB839 (yellow) binds the GLS dimer-dimer interface (one GLS monomer is colored blue) using residues from the activation loop. The sites of the disease-associated mutations S482C and H461L are shown in pink. Image generated from PDB ID 5JYO using the USCF ChimeraX software.[51]
